# Replicability of bulk RNA-Seq differential expression and enrichment analysis results in cancer research

**DOI:** 10.1101/2023.10.25.563901

**Authors:** Peter Degen, Matúš Medo

## Abstract

The high-dimensional and heterogeneous nature of transcriptomics data from RNA sequencing (RNA-Seq) experiments poses a challenge to routine down-stream analysis steps, such as differential expression analysis and enrichment analysis. Additionally, due to practical and financial constraints, RNA-Seq experiments are often limited to a small number of biological replicates; three replicates is a commonly employed minimum cohort size. In light of recent studies on the low replicability of preclinical cancer research, it is essential to understand how the combination of population heterogeneity and underpowered cohort sizes affects the replicability of RNA-Seq research. Using 7’000 simulated RNA-Seq experiments based on real gene expression data from seven different cancer types, we find that the analysis results from underpowered experiments exhibit inflated effect sizes and are unlikely to replicate well. However, the ground-truth results obtained by analyzing large cohorts show that the precision of differentially expressed genes can be high even for small cohort sizes. The poor replicability of underpowered experiments is thus a direct consequence of their low recall (sensitivity). In other words, the low replicability of underpowered RNA-Seq cancer studies does not necessarily indicate a high prevalence of false positives. Instead, the results obtained from such studies are limited to small and mostly random subsets of a larger ground truth. We conclude with a set of practical recommendations to alleviate problems with underpowered RNA-Seq studies.

**Author Summary:** Transcriptomics data from RNA sequencing (RNA-Seq) experiments are complex and challenging to analyze due to their high dimensionality and variability. These experiments often involve limited biological replicates due to practical and financial constraints. Recent concerns about the replicability of cancer research highlight the need to explore how this combination of limited cohort sizes and population heterogeneity impacts the reliability of RNA-Seq studies. To investigate these issues, we conducted 7’000 simulated RNA-Seq experiments based on real gene expression data from seven different cancer types. We show that experiments with small cohort sizes tend to produce results with exaggerated effects that can be difficult to replicate. We further found that while underpowered studies with few replicates indeed lead to little-replicable results, the identified differentially expressed genes are reliable as shown by low rates of false positives. Each underpowered study thus discovers a small subset of the ground truth. Our study concludes with practical recommendations for RNA-Seq studies with small cohort sizes.

## 1 Introduction

The rapidly increasing availability of large and highly heterogeneous omics data from high-throughput sequencing technologies has stimulated the development of appropriate statistical methods [1]. Differential expression analysis, the problem of detecting systematic differences in the expression levels of genomic features between experimental conditions (e.g., normal tissue versus tumor tissue), is a key problem in this field [2–4]. The term “genomic features” here can refer to genes, exons, transcripts, or any other genomic region of interest; we shall henceforth use the umbrella term “gene” for simplicity of notation. Gene expression levels are typically quantified using read counts obtained from next-generation sequencing technologies such as RNA-Seq [5]. Due to various sources of biological and technical variability, statistical hypothesis tests are needed to determine the significance of any observed difference in read counts. Genes that pass a significance threshold after correcting for multiple hypothesis testing are designated as differentially expressed genes (DEGs). They can be used for further downstream analysis such as enrichment analysis [6] or independent scrutiny in subsequent wet lab experiments.

The statistical power of RNA-Seq experiments naturally increases with the number of patients (biological replicates). However, a review of the available literature suggests that actual cohort sizes often fall short of the recommended minimum cohort sizes. For example, Schurch et al. [7] estimated that at least six biological replicates per condition are necessary for robust detection of DEGs, increasing to at least twelve replicates when it is important to identify the majority of DEGs for all fold changes. Lamarre et al. [8] argued that the optimal FDR threshold for a given replication number *n* is 2^*−n*^, which implies 5–7 replicates for typical thresholds of 0.05 and 0.01. Bacarella et al. [9] cautioned against using fewer than seven replicates per group, reporting high heterogeneity between the analysis results depending on the choice of the differential expression analysis tool. Ching et al. [10] estimated optimal cohort sizes for a given budget constraint, taking into account the trade-off between cohort size and sequencing depth. While emphasizing that the relationship between cohort size and statistical power is highly dependent on the data set, their results suggest that around ten replicates are needed to achieve ≳ 80% statistical power.

Despite this body of research warning against relying on insufficient replication numbers, three replicates per condition remains a commonly used cohort size, and most RNA-Seq experiments employ fewer than five replicates [7]. A survey by Baccarella et al. [9] reports that about 50% of 100 randomly selected RNA-Seq experiments with human samples fall at or below six replicates per condition, with this ratio growing to 90% for non-human samples. This tendency toward small cohorts is due to considerable financial and practical constraints that inhibit the acquisition of large cohorts for RNA-Seq experiments, as including more patients in a study requires substantial time and effort, especially for rare disease types. In light of this discrepancy between the recommended and actual cohort sizes, more research is urgently needed on the potentially detrimental effects of low-powered RNA-Seq experiments. Unfortunately, the recent literature on this subject is limited.

One recent study was conducted by Cui et al. [11], who subsampled RNA-Seq data from The Cancer Genome Atlas (TCGA) and calculated the overlap of DEGs among the subsampled cohorts. Noting the low overlap of results for small cohort sizes, the authors recommend using at least ten replicates per condition and interpreting low-powered studies with caution. Another study based on the TCGA data was conducted by Wang et al. [12], whose primary concern was the comparison of different metrics for evaluating replicability. The authors report significantly heterogeneous results depending on the chosen replicability metric and the studied cancer type.

More generally, Ioannidis [13] proposed a simple statistical model for high-throughput discovery-oriented research, of which RNA-Seq is a prime example. This model can be used to demonstrate potentially high rates of false positive results. Although the author’s claim that “most published research findings are false” has been the subject of considerable debate [14, 15], it indeed appears to be the case that certain fields such as preclinical cancer biology are struggling with a high prevalence of research with replication problems [16, 17]. Errington et al. [18] recently conducted a large-scale replication project that attempted to replicate 158 effects from 50 experiments across 23 high-impact papers in preclinical cancer research, achieving a success rate of 46%. Furthermore, the authors found that 92% of the replicated effect sizes were smaller than in the original study.

Motivated by the described issues, our goal is to investigate the replicability of RNA-Seq analysis results in the context of preclinical cancer research. Compared to the studies by Cui et al. [11] and Wang et al. [12], we comprehensively explore the space of various analysis parameters and decisions, including cancer type, cohort size, choice of the differential expression analysis tool, choice of the significance threshold, filtering differentially expressed genes by their fold change, outlier removal, and downstream enrichment analysis methods. This extensive experimentation allows us to formulate a set of recommendations for researchers working in this domain.

## 2 Results

### 2.1 Low DEG replicability for small cohort sizes

We begin by exploring the dependencies of the DEG replicability on the cohort size. Fig. 1 shows the median number of DEGs and the corresponding median DEG replicability as a function of the cohort size for different statistical tests and fold change thresholds, aggregated across all seven studied cancer types. The crosses in the bottom row depict the expected replicability under the null model of random gene selection given in Eq. (3), using the median number of identified DEGs for both *S*_*i*_ and *S*_*j*_.

**Fig. 1.**
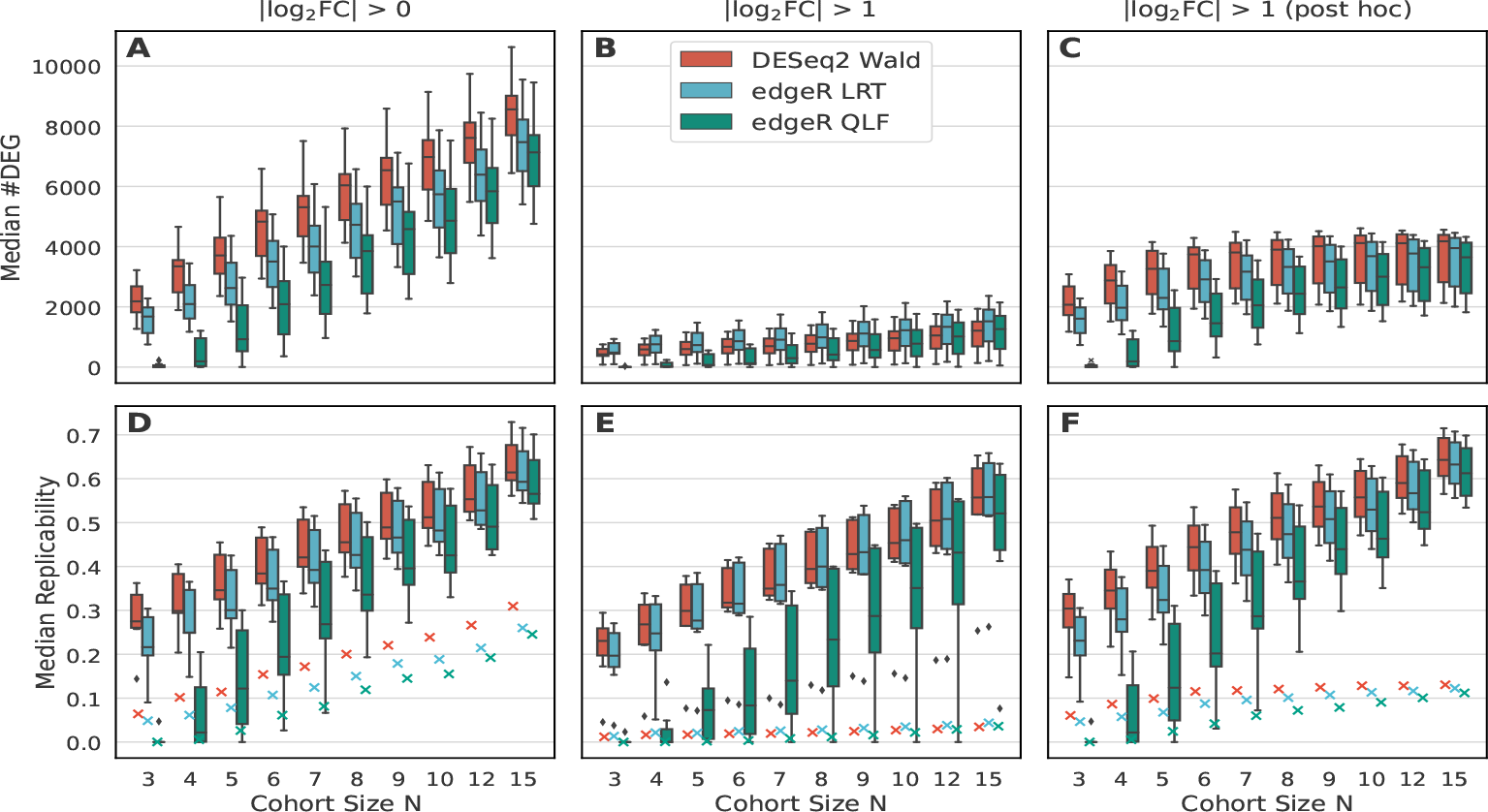
Dependencies on the cohort size: median number of DEGs (**A-C**) and median DEG replicability (**D-F**). Each box plot summarizes seven data points that represent the median values obtained using 100 simulated cohorts for one of the seven cancer types studied. In the bottom row, the replicability values expected for randomly selected DEGs are depicted as crosses. Left column: DEGs defined without a fold change threshold. Middle column: DEGs defined using built-in statistical methods that test for the threshold |log_2_ FC| *>* 1. Right column: DEGs as in the first column with an additional threshold on the estimated fold changes.

Regardless of the method, cancer type, and fold change threshold, we observe consistently low DEG replicability for small cohort sizes, only reaching ≈ 50% for *N* ≳ 10. Without a fold change threshold (left column), the Wald test from *DESeq2* shows consistently higher replicability compared to both *edgeR* methods (LRT and QLF) and also declares more genes as differentially expressed. QLF is the most conservative method, resulting in the fewest DEGs and, in turn, the lowest replicability. The same trends are observed when the DEGs are filtered with a post hoc fold change threshold (right column). However, when formally testing for a minimum significant fold change (middle column), Wald and LRT perform comparably.

We also observe that imposing a fold change threshold (formally or post hoc) has only a minor impact on replicability. However, replicability alone is not enough to assess the results. When the number of DEGs is high, a sizeable part of replicability is due to the expected replicability under the null model (the crosses in Figure 1). If we judge the results by the difference between the measured and expected replicability, it becomes clear that using a fold change threshold is beneficial.

Supplementary Fig. 3 in Additional File 1 shows replicability for individual cancer types. The most striking cancer type is prostate adenocarcinoma (PRAD), which has the lowest replicability (e.g., all outliers marked in panel E of Fig. 1 correspond to PRAD). This cancer type also has the lowest number of ground truth DEGs when the fold change is controlled formally (around 500 DEGs compared to 1200–3000 for the other cancer types; see Supplementary Fig. 1 in Additional File 1).

### 2.2 Ground truth comparison

The empirical ground truth of DEGs is obtained using all patients in a given data set and taking the intersection of the results obtained using all three methods for differential expression analysis. Depending on the data set, the ground truth comprises between 9’585 and 13’735 DEGs with | log_2_ FC| *>* 0 and between 534 and 3’024 DEGs with | log_2_ FC| *>* 1 (Supplementary Fig. 1).

The top row of Fig. 2 shows the precision of DEGs as a function of the cohort size. At the FDR threshold of 5%, the expected precision is 1 − 5% = 0.95, represented as a solid black line. Not depicted in the figure is the expected precision for the null model of random gene selection, which is 70%, 11%, and 19% for panels A, B, and C, respectively, averaged over all cancer types and differential expression methods.

**Fig. 2.**
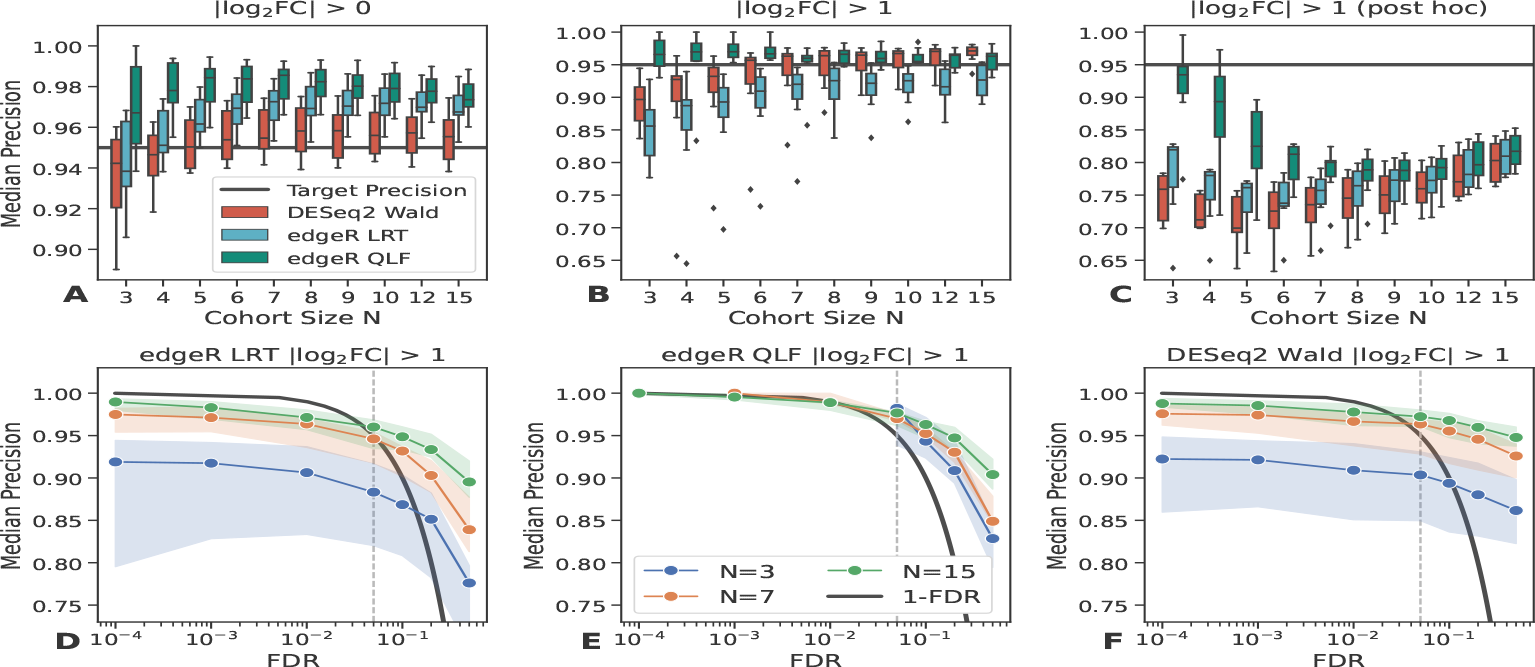
**A-C**: Precision of DEGs as a function of the cohort size. The target precision of 1-FDR = 95% is depicted as a solid black line. All outlier points (black diamonds) originate from the PRAD data set. For panel B and *N* = 3, one outlier point for each box plot falls outside the figure range (the values are 57% (Wald), 53% (LRT), and 13% (QLF)). **D-F**: Precision as a function of the FDR threshold for DEGs defined using a formal fold change threshold. Shown is the median value across the seven cancer types plus 95% confidence interval bands. The vertical dashed line corresponds to the FDR threshold of 5% used in the top row. The solid black line corresponds to the ideal case where the empirical FDR is the same as the chosen FDR threshold. In panel E, results are not shown for small cohort sizes and stringent FDR thresholds as the conservative edgeR QLF test then does not detect enough DEGs for a meaningful evaluation of precision.

In panel A, when no fold change threshold is applied, we see that the target precision of 95% is best achieved by the Wald test from *DESeq2*, whereas both *edgeR* tests (QLF and LRT) exceed the expected precision by 2–3%. In panel B, we statistically test log_2_ FC *>* 1. For small cohorts *N* ≲ 6, we observe that Wald and LRT fail to control the precision at 95%, while QLF shows the most robust control across the entire range of *N*.

In panel C, we test the null hypothesis of zero fold change and subsequently filter the significant genes that have |log_2_ FC| *>* 1. Because of this post hoc filtering, the precision is markedly lower than the target precision expected from FDR thresholding; it is thus better to avoid this filtering approach. The low precision values are a direct consequence of inflated fold change estimates in small cohorts (see Fig. 5D, E below): Many genes that pass the fold change threshold for *N* = 5, for example, are false positives as their ground truth fold change values are substantially smaller.

The bottom row of Fig. 2 shows the precision of DEGs as a function of the FDR threshold for the (formal) threshold log_2_ FC *>* 1. Also depicted as a black line is the ideal case where the empirical precision is the same as one minus the chosen FDR threshold (the target precision). For stringent thresholds ≲ 1%, we observe that the Wald and LRT tests do not reach the target precision even for *N* = 15. In comparison, the QLF test manages to converge on the target precision as long as there is sufficient power to detect any DEGs at all. At stringent FDR thresholds, this is only possible with large cohorts due to the conservative nature of the test.

Fig. 3 depicts the MCC and recall metrics as a function of the cohort size. In contrast to precision, these two metrics are strongly correlated with the cohort size, regardless of the thresholding of fold change. In the left column, when no threshold is applied, we observe that Wald is the best-performing method. The high precision observed earlier for QLF and LRT results here in a comparatively lower recall. In fact, *edgeR* recall is so low that both tests are consistently outperformed by the Wald test with respect to the MCC metric that balances all four categories of the confusion matrix. The same is true when we apply a post hoc fold change threshold (right column). However, when we statistically test | log_2_ FC| *>* 1 (middle column), the trend between Wald and LRT is reversed across the entire range of *N*. Again, this is consistent with the precision trends observed earlier. Regardless of the method, applying a fold change threshold substantially improves the MCC compared to not using a threshold.

**Fig. 3.**
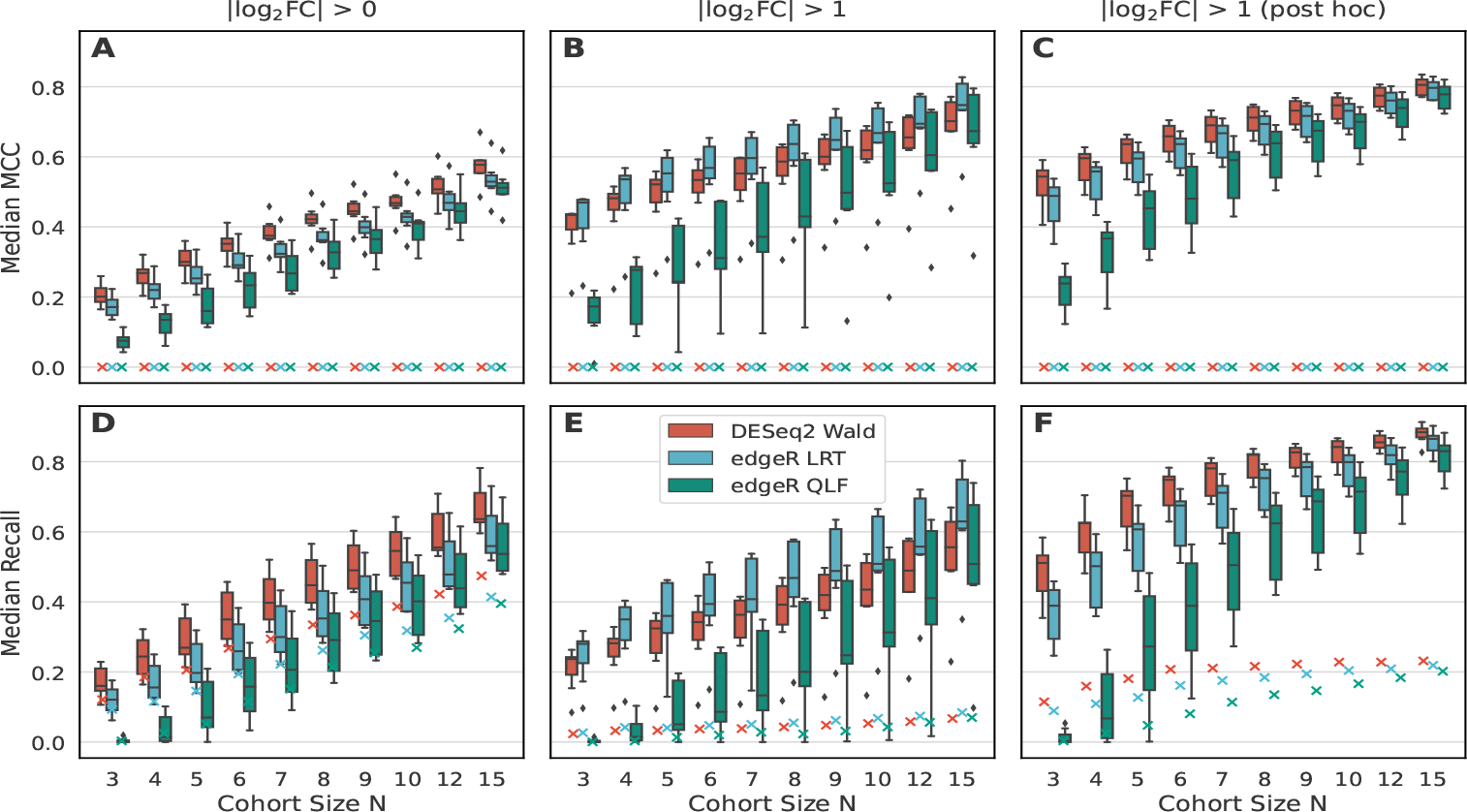
MCC and recall of DEGs as a function of the cohort size. The expected metrics for the null model of random gene selection are shown as crosses.

### 2.3 Low enrichment replicability for small cohort sizes

We now use the output of individual methods for differential expression analysis for a common downstream step: gene enrichment analysis. This allows us to compute the replicability of enriched GO terms and KEGG pathways for different enrichment analysis methods and cohort sizes.

In Fig. 4, the replicability as a function of the cohort size is depicted for the GO terms (top row) and KEGG pathways (bottom row). The first column depicts the results from the GSEA enrichment analysis method for two different ranking metrics, whereas the remaining columns show the results from over-representation analysis (ORA) for different DEG methods and fold change thresholds. Although the replicability shows substantial differences depending on the chosen method, in general, we can say that the replicability is low for small cohorts and increases with the cohort size. For the GO terms, the highest replicability is achieved with *DESeq2* Wald and a post hoc fold change cutoff. For the KEGG pathways, the highest replicability is achieved with GSEA and log fold change ranking.

**Fig. 4.**
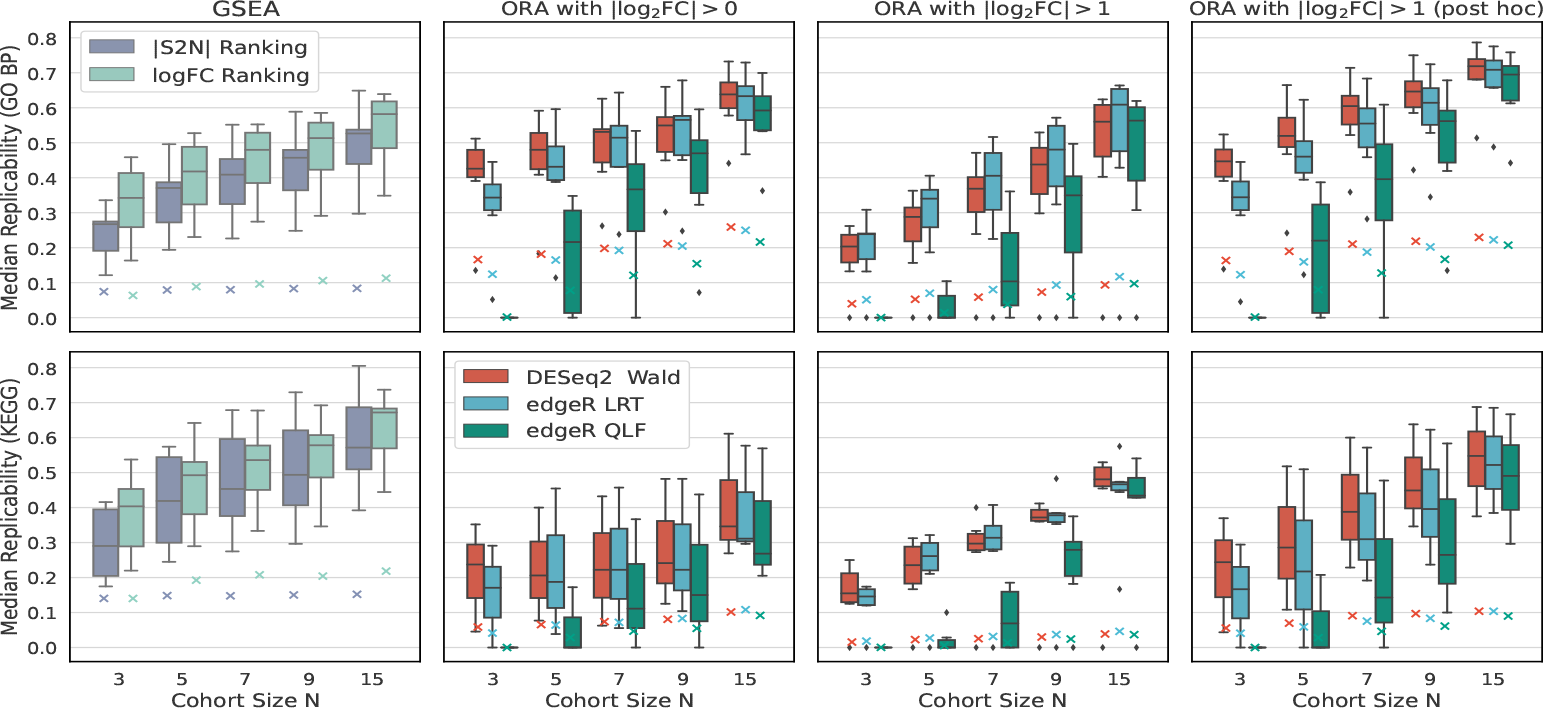
Replicability for enriched GO terms (top row) and KEGG pathways (bottom row). Each box plot summarizes seven data points, representing the median values in 100 cohorts for each cancer type.

**Fig. 5.**
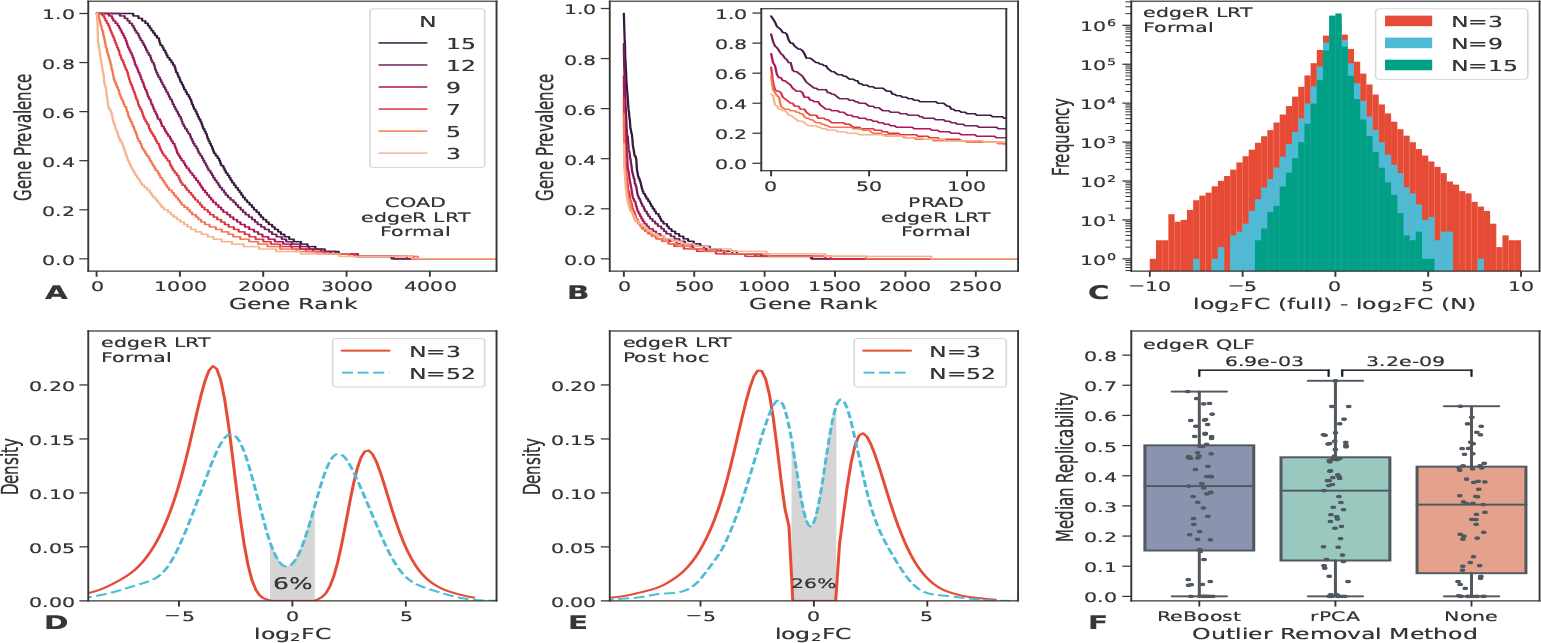
**A-B**: Gene prevalence when using *edgeR* LRT with a formal fold change cutoff for two datasets and different cohort sizes. **C**: Genome-wide fold change estimates obtained from the full data sets (median *N* = 52) of all cancer types minus fold change estimates obtained from small cohorts (*N* = 3, 9, 15). **D**: Density of fold change estimates for DEGs obtained from small cohorts with *N* = 3 (solid line) and density of fold changes for the same genes using the full data sets for fold change estimation (dashed line). The gray-shaded area represents the fraction of genes that spuriously passed the threshold for *N* = 3 but would not have passed in a more powerful study. DEGs were identified with *edgeR* LRT and a formal fold change threshold. **E**: Same as panel D, but using a post hoc fold change threshold instead. **F**: Replicability of DEGs identified with *edgeR* QLF (without fold change threshold) for various outlier removal methods. Each symbol represents one cancer type and cohort size. A Wilcoxon signed-rank test with paired samples was used to obtain the p-values above the box plots.

Additionally, Figs. B5 to B8 in Additional file 1 show analogous figures for the MCC, precision, recall, and the number of significant terms and pathways. In particular, for ORA with a formal (post-hoc) fold change threshold, the precision of the enriched GO terms is fairly stable at ≈ 88% (≈ 85%) for all methods and cohort sizes. The precision of KEGG terms is ≈ 14% lower, but also relatively stable for different cohort sizes. However, it should be noted that all these results are still considerably lower than the expected precision of 1 − FDR = 95%.

### 2.4 Differential expression prevalence

Upon observing a low replicability value, it is still possible that some genes are differentially expressed in (almost) all simulated cohorts or that all genes are subject to the same level of variability. To clarify these two possibilities, we devise a new metric, the differential expression prevalence, which is defined as the fraction of cohorts in which a given gene is identified as differentially expressed. We compute the gene prevalence for *edgeR* LRT with a fold change cutoff. Fig. 5A shows results for the COAD data set; these results are representative of most analyzed cancer types. While there are only 13 genes that have DE-prevalence 1 (i.e., they are always differentially expressed) at *N* = 3, this number grows to 756 for *N* = 15. Prostate adenocarcinoma (Fig. 5B) once again exhibits abnormal behavior as there are no genes with DE-prevalence 1 for cohorts as large as 12. This can be a sign of the disease heterogeneity that we have not managed to suppress by the described process of RNA-Seq data acquisition and cleaning. Supplementary Figs. 17-18 in Additional file 1 show the gene prevalence for all cancer types and differential expression methods.

### 2.5 Inflated effect sizes for underpowered studies

The potential of underpowered (lacking sufficient sample size) studies to inflate effect sizes and undermine the reliability of results has been reported in a range of fields [19– 21]. We demonstrate this effect for RNA-Seq data using aggregated results from all subsampled cohorts and cancer types. To this end, we define a ground truth for the fold change estimates by running *edgeR* on the entire data sets. As before, the choice of the analysis tool is unimportant because the fold change estimates are similar between the tools.

Fig. 5C depicts the frequency of fold change deviations for all genes and three distinct cohort sizes, relative to the ground truth. With larger cohorts, the width of the distribution becomes substantially narrower, demonstrating the importance of sample size for reliable fold change estimation. In Fig. 5D, we narrow our focus from all genes to DEGs with |log_2_| *>* 1 and FDR *<* 0.01 obtained from small cohorts (*N* = 3) with *edgeR* LRT. The solid red line depicts the kernel density of fold change estimates for the identified DEGs, where the fold changes have been estimated directly from these small cohorts. The blue dashed line represents the fold change density for exactly the same DEGs, but using the entire data sets for the fold change estimation (median *N* = 52). Here, the inflation of effect sizes is clearly visible in the form of a systematic bias towards large fold change estimates for the small cohorts. Around 6% of the genes (gray area) that passed the fold change threshold with *N* = 3 would not have passed the threshold if the fold changes had been estimated with the entire data set. When using a post hoc fold change threshold, this number increases to 26% (Fig. 5E).

### 2.6 Removing outlier patients

An outlier patient whose differential expression pattern is different from that of the majority of samples in a cohort can hamper the identification of differentially expressed genes. As we have seen that low recall is the main cause of low replicability of DEGs, we now examine whether it is possible to counteract this issue by removing potential outlier patients. Our *ReBoost* outlier removal method is based on removing matching samples corresponding to one or more patients so that the number of identified DEGs is maximized (see Methods for details). In addition to the newly introduced method, we also tested the recently proposed rPCA method [22] for outlier removal. We have tested also a third tool, the molecular degree of perturbation (MDP) [23]. However, we found that MDP was able to detect outliers only in a small fraction of simulated cohorts, its performance is thus similar to not removing any outliers at all (the difference is not statistically significant). We decided to save computational resources by not applying MDP to all 7’000 cohorts.

Fig. 5F shows the replicability of DEGs with and without the removal of outlier patients (no fold change thresholds are used here). Aggregated over all cancer types and cohort sizes, we find that outlier removal yields significant replicability increases that are particularly large when edgeR QLF is used for differential expression analysis.

Supplementary Figs. 9, 10 and 12 in Additional file 1 show analogous plots for the replicability, MCC, precision, and recall, with and without post hoc fold change cutoff. As the *ReBoost* method is designed to increase recall at a fixed FDR threshold, it shows the highest increase in recall in all tested scenarios. Unsurprisingly, this results in a small precision loss. As we have already shown that QLF and LRT exceed the target precision of 1 − *FDR*, this decrease in precision can be tolerated. If no fold change thresholding is used, the *ReBoost* method also shows the highest improvement in MCC across the three tested differential expression methods. Supplementary Figs. 13 to 16 in Additional file 1 show the impact of outlier removal for individual cancer types and cohort sizes. The removal of outliers produces the biggest improvements in MCC and replicability for THCA and PRAD—two cancer types whose replicability is particularly low for small cohort sizes.

## 3 Discussion

We have comprehensively analyzed the replicability of the results of RNA-Seq differential expression analysis. Compared to recent work by Cui et al. [11], we used data from more cancer types, considered multiple tools for the analysis, characterized fold change thresholding strategies and the impact of outlier removal, and evaluated the impact of low replicability on downstream enrichment analysis (see Table A2 in Additional file 1 for a detailed comparison of the methodology used). We support their conclusion that differential expression results obtained from small cohorts (*N* ≲ 10) lead to low inter-experiment replicability (they use the term “overlap rate”). Contrary to [11], our results show that low replicability does not imply that the DEGs do not generalize to larger cohorts. This is demonstrated by the median precision above 95% in Figure 2 even for the smallest tested cohort size of *N* = 3 (for DESeq2 Wald and edgeR QLF without a fold change cutoff).

We find instead that low replicability values (Figure 1) are a direct consequence of low recall and frequent false negatives (Figure 3). Therefore, a possible failure to replicate a low-powered RNA-Seq study indicates that the two studies are looking at different small parts of a larger ground truth (Fig. 6).

**Fig. 6.**
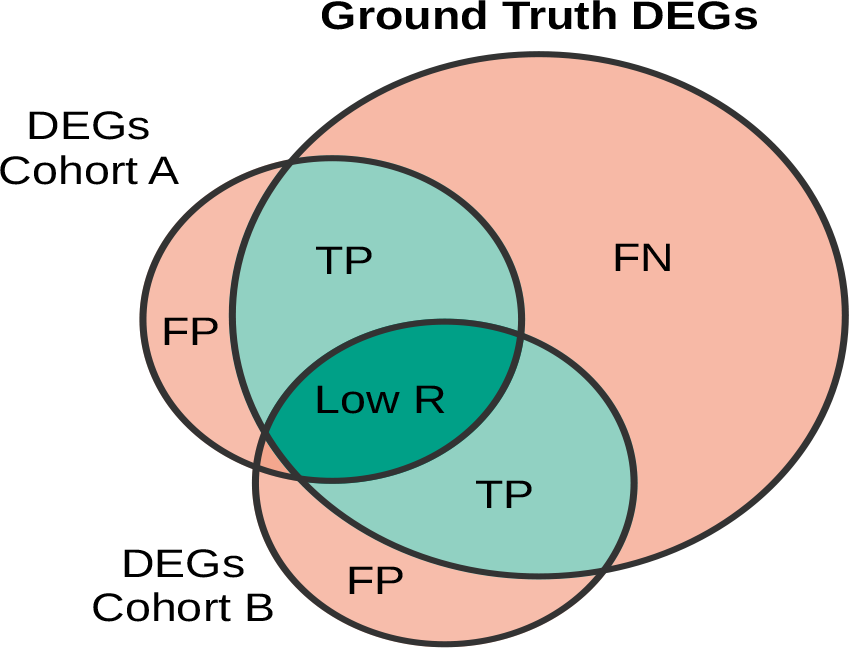
An illustrative Venn diagram that summarizes our results. DEGs obtained by analyzing small cohorts A and B typically have little overlap (dark green area), implying low replicability. At the same time, their overlap with the ground truth DEGs obtained by analyzing a much bigger cohort is large (dark and pale green areas) and the fraction of false positives (FP) is small. This is made possible by a large number of false negatives (FN).

Li et al. [24] recently reported that edgeR and DESeq2 suffer from false positives, as demonstrated by the high number of DEGs they identify in permuted input datasets where significant differences are assumed to be removed by permutations. This issue does not seem to be relevant for the RNA-Seq data analyzed here as: (1) DEGs start to appear in permuted data only when the number of samples is large (*n* ≳ 32) and (2) the numbers of DEGs thus found are much smaller compared to the numbers of 15 DEGs found in the unpermuted data (Supplementary Fig. 19 in Additional file 1). In [24], the most striking observations have been made for an immunotherapy study. We have noticed that some highly expressed genes in this dataset have several zero counts that are known to cause problems for *edgeR* [25]; this could have contributed to the reported behavior.

We confirm the results of Chen et al. [22] that removing outliers can substantially increase the recall of DEGs. In the case of our custom-made *ReBoost* method, recall gain outweighs precision loss to such a degree that the MCC metric is improved as well. Additionally, replicability is also improved, as low recall is the main driver of low replicability in underpowered RNA-Seq studies. Nevertheless, removing outliers from experimental data remains a contested topic, with some researchers even encouraging systematic heterogenization of study samples [26]. We recommend manually inspecting the identified outliers to assess their biological importance. For larger cohorts, detection of patient subpopulations could be considered as an alternative to outlier removal.

Moving on from significant DEGs to enriched gene sets (GO terms and KEGG pathways), we observe a similar pattern of low replicability and recall for small cohorts. The prevalence of false positives is more problematic here: In all scenarios tested, the target precision of 1 − FDR = 95% was not reached (Fig. 6 in Supplementary File 1). The precision of DEGs is particularly low for overrepresentation analysis (ORA) without a fold change threshold, which is thus to be avoided. In the best-case scenario, ORA with a fold change threshold comes within *<* 10% of the target precision for GO terms. Overall, these results suggest that enrichment analysis is too sensitive to perturbations in input data to faithfully reproduce findings obtained from cohorts with vastly different sizes.

We next move to the question of a minimum recommended cohort size. Inspecting the precision of DEGs in Fig. 2, we suggest that cohorts with fewer than 6 replicates per condition should be avoided due to the precision below 1 − *FDR* for 2 out of 3 differential expression tests (LRT and Wald test). Although the QLF test manages to achieve high precision even for *N <* 6, it often fails to detect any significant genes at such small cohort sizes. Beyond the lower limit of 6 replicates, we observe gradual improvements in recall and replicability for all three evaluated methods, with slightly reduced improvements for *N* ≳ 10. These numbers are in general agreement with previous recommendations on cohort sizes [7–11].

We proceed with a discussion of the limitations of our study. Firstly, our study is based on relatively small source data sets (median *N* = 52). This means that our subsampled cohorts inevitably contain repeating patients. The most extreme case is given by the largest subsampled cohort size of *N* = 15 for the smallest data set, COAD, with 39 replicates. In this case, there are 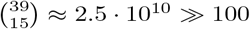 distinct subsampled cohorts. However, the expected number of patients shared between two subsampled cohorts is 15^2^*/*39 ≈ 6. These repeated patients lead to an underestimation of population heterogeneity and thus our replicability estimates can be considered conservative in this respect. One way to avoid repeated patients would be to resample pairs of *N* patients from the source data without replacement. Using the same computational budget of 100 analyses (per cancer type per cohort size), this approach would yield 50 pairs of cohorts and 50 replicability values. To get more statistics, we chose to resample with replacement and then calculate the replicability between all pairs of 100 cohorts, yielding 4’950 replicability values.

Another limitation is our focus on bulk RNA-Seq cancer data. Generalizability of our results to other research domains (with possibly less heterogeneous populations than cancer) and newer technologies such as single-cell RNA-Seq remains to be tested. However, due to the substantial increase in complexity and computational requirements of single-cell analysis pipelines, it could be challenging to perform an analogous single-cell replicability study that relies on large numbers of subsampled cohorts. We instead highlight a recent relevant study by Squair et al. [27], who showed that statistical methods for differential expression analysis that do not account for biological variability are prone to false discoveries in single-cell data.

As a side product of our investigation, we conclude by briefly discussing our GSEA results in the context of a study by Zyla et al. [28], who tested the robustness of different gene ranking methods for GSEA. Contrary to their results, we find that ranking the genes by their log_2_ FC yields better performance metrics (MCC and replicability) than ranking them by the absolute signal-to-noise ratio. However, it should be noted that there are several methodological differences between our studies that could be responsible for this discrepancy. Notably, we used gene permutations to calculate the p-values of enriched gene sets, whereas Zyla et al. used sample permutations to preserve the gene correlation structure in the data. In our case, sample permutations were not feasible because we were explicitly interested in small cohort sizes (the GSEApy package recommends sample permutations only if the number of samples is ≥ 15). Other methodological differences in Zyla et al. include the exclusive use of unpaired study designs and a different procedure to correct for multiple hypothesis testing. When compounded with the differences in datasets used, we see no immediate cause for concern in the discrepancy of results.

## 4 Conclusion

Although the broader replication crisis in science [16–18, 29] includes numerous external, human factors such as dysfunctional incentive systems, selective reporting, inadequate statistical training, and publication bias, here, we assumed otherwise ideal research practices and only concerned ourselves with low replicability arising from underpowered studies of heterogeneous biological populations. Crucially, we showed that the low replicability of underpowered RNA-Seq studies need not be a cause for low confidence in the original findings, as the low replicability is driven mainly by false negatives and not by false positives. However, we also found that there is a systematic inflation of effect sizes for underpowered RNA-Seq studies (Fig. 5D). To summarize these findings for biologists working with RNA-Seq data from small cohorts sampled from a heterogeneous population:

- Significant DEGs from one small cohort are unlikely to be significant in another small cohort. At the same time, they are likely to be significant in a much larger cohort.
- The magnitudes of significant fold changes observed on the transcriptome level are likely to decrease in subsequent validation attempts, e.g., on the proteome level.
- We recommend a minimum cohort size of 6 biological replicates per condition.
- *DESeq2* Wald with fold change cutoff performed best in our evaluation: Its precision is close to the expected value and recall is high. For very small cohorts (*N* = 3, 4), *edgeR* QLF yields better precision, but most likely few to no DEGs are found.
- If lower recall is acceptable when performing enrichment analysis, ORA with a fold change cutoff based on *edgeR* LRT (or *DESeq2* Wald) is the best option due to precision close to 90% for most cancer types and cohort sizes. Otherwise, GSEA based on the fold change ranking provides higher recall (and lower precision).

## 5 Methods

Our main strategy to investigate the replicability of RNA-Seq analysis results is based on repeatedly subsampling small cohorts from large data sets and determining the level of agreement between the analysis results based on these subsamples (Fig. 7). To this end, we used The Cancer Genome Atlas (TCGA) public data repository to obtain RNA-Seq data sets with enough patients. For improved statistical power [30], we limited our study to paired design experiments with one normal tissue sample and one matching primary tumor sample for each patient. As listed in Table 1 and further described in the next section, we identified seven data sets from different types of cancer that matched our criteria. For each cancer type and target cohort size *N* = {3, 4, 5, 6, 7, 8, 9, 10, 12, 15}, we simulated 100 RNA-Seq experiments by randomly selecting a small cohort of size *N* from the full data set of the given cancer type. These simulated experiments can be thought of as independent studies aiming to answer the same research question using the exact same methods but based on different cohorts drawn from the overall population. Each simulated cohort consists of unique patients, but a patient may appear in multiple cohorts. In total, we simulated 7’000 RNA-Seq experiments and analyzed each one using multiple analytical pipelines.

**Table 1.** The number of genes after filtering of lowly expressed genes with *edgeR* and the number of patients (biological replicates) for each of the seven data sets (cancer types) considered in this paper. Each patient contributes one normal tissue sample and one primary tumor sample to the data set.

| Data set | TCGA project name | Genes | Patients |
| --- | --- | --- | --- |
| BRCA | Breast invasive carcinoma | 18'737 | 111 |
| KIRC | Kidney renal clear cell carcinoma | 18'231 | 72 |
| THCA | Thyroid carcinoma | 18'952 | 58 |
| LUAD | Lung adenocarcinoma | 17'681 | 52 |
| PRAD | Prostate adenocarcinoma | 18'880 | 51 |
| LIHC | Liver hepatocellular carcinoma | 15'607 | 50 |
| COAD | Colon adenocarcinoma | 17'220 | 39 |

**Fig. 7.**
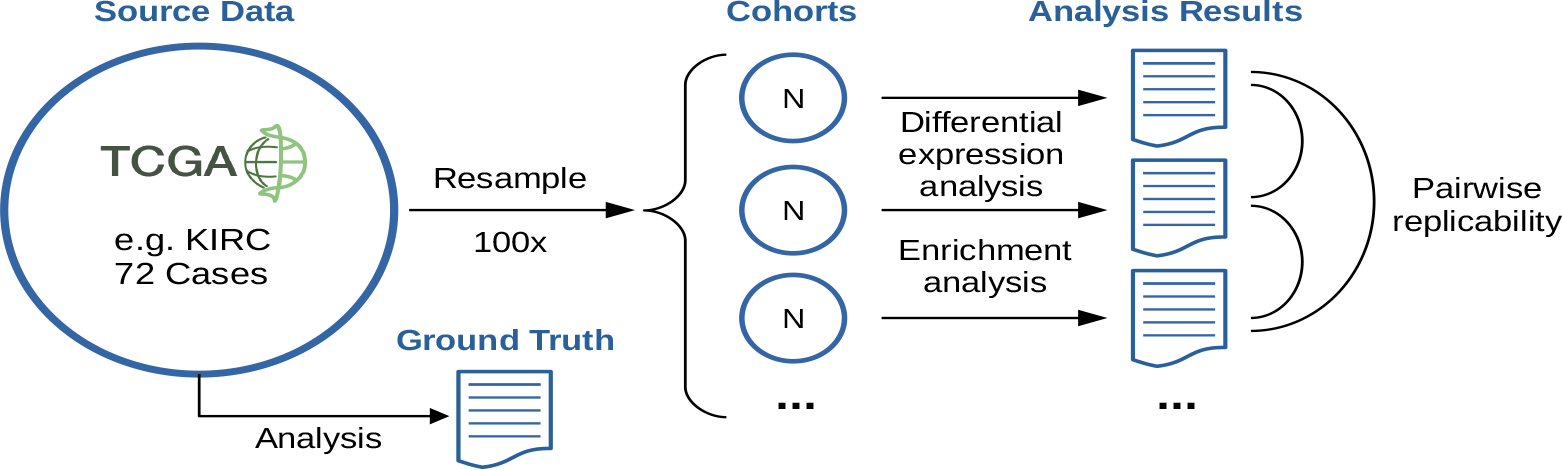
Flowchart of our study design. An RNA-Seq data set obtained from the TCGA is subsampled to yield 100 small cohorts of size *N*, which are analyzed separately and compared pairwise to measure the level of agreement between the results. Additionally, the same analysis steps are run on the entire source dataset to define our ground truth. This procedure is repeated for seven different cancer types and ten different cohort sizes ranging from 3 to 15 (7’000 cohorts in total).

### 5.1 Data

Table 1 lists the seven data sets used in this study. All data sets consist of unnormalized integer read counts that have already been pre-processed and aggregated to quantify gene expression levels. We downloaded the data sets using a custom Python script that accessed the API of the Genomic Data Commons (GDC) of the United States National Cancer Institute [31]. For each primary cancer site, we filtered the available cases by experimental strategy (RNA-Seq) and data category (transcriptome profiling). We excluded all patients who did not have at least one primary tumor tissue sample and one solid normal tissue sample. If the cases came from multiple projects, we kept the patients from the most populated project. There were seven projects with at least 50 patients. To avoid excessive cohort heterogeneity, we finally kept only patients with the most common disease type for the given project. The median number of patients across the seven resulting data sets is 52 (range 39–111). Finally, for each data set, we filtered lowly expressed (and hence uninformative) genes using the *filterByExpr* function from *edegR*

### 5.2 Differential expression analysis

For each simulated experiment, we determined the genes that are differentially expressed between normal and primary tumor tissue samples using the popular R packages *edgeR* [4] and *DESeq2* [3]. Both packages rank among the leading tools when considering small sample sizes [7] and overall performance [10], boasting 31’1110 and 51’123 respective citations on Google Scholar as of June 15, 2023.

Before testing for differential expression, we normalized the counts using the *calcNormFactors* function from *edgeR* and *estimateSizeFactors* from *DESeq2*, respectively. In both cases, a paired-sample design matrix was used to improve statistical power [10]. Unless otherwise noted, we determined the significant DEGs using a 5% threshold on the Benjamini-Hochberg adjusted p-values to control the false discovery rate (FDR); we will henceforth use the term “FDR” to refer to the adjusted p-values. To test for differential expression, we considered several statistical approaches, as listed in Table 1 in Additional file 1 and described in the next two paragraphs.

First, an important issue that needs to be addressed is the question of a minimum absolute fold change, below which genes are not considered to be biologically relevant. There are two approaches to filtering genes with small fold change. The statistically principled way is to formally test the null hypothesis |log_2_ FC| ≤ *t*_*null*_, where *t*_*null*_ is the chosen minimum significant threshold [32]. However, many practitioners instead use an alternative approach that we will designate as post hoc thresholding (also called double filtering in [33]). In this approach, only the null hypothesis of a zero fold change is formally tested, followed by a filtering step in which the significant genes whose estimated fold change is below a chosen cutoff are removed. One disadvantage of this approach is that the FDR is no longer properly controlled at the specified significance level, as we will see in the results section. Compared to a formal statistical test, post hoc thresholding is also a more permissive approach, as genes that barely pass the threshold might not reach statistical significance in a formal test but are still retained when using a post hoc cutoff. For comparison, we include both types of fold change thresholding in our analysis.

For *DESeq2*, we used the default Wald test, which tests for differential expression above a user-specified absolute log_2_ fold change threshold. In our case, we chose two thresholds, *t*_*null*_ ∈ {0, 1}. For *edgeR* and *t*_*null*_ = 0, we used both the likelihood-ratio test (LRT) and the quasi-likelihood F-test (QLF). QLF is described by the authors as offering more conservative and reliable type I error control when the number of replicates is small [34]. For *t*_*null*_ = 1, we used the t-test relative to a threshold (TREAT) [32], as recommended by the authors. Like LRT and QLF, TREAT is a parametric method that requires negative binomial models to be fitted to the data before any testing can be done. Users can choose between the functions *glmFit* or *glmQLFit*, which are the same functions used for the LRT and QLF pipelines, respectively. For this reason, we keep the labels LRT and QLF to designate our use of TREAT.

### 5.3 Enrichment analysis

We performed enrichment analysis for each simulated experiment to investigate the impact of small cohort sizes on the results of the downstream analysis. We considered two commonly used types of enrichment analysis: Over-representation analysis (ORA) and Gene Set Enrichment Analysis (GSEA) [6]. In both cases, we tested for significantly enriched pathways from the Kyoto Encyclopedia of Genes and Genomes (KEGG), as well as enriched Gene Ontology (GO) terms from the Biological Process subdomain (BP).

The first method, ORA, was applied using the R package *clusterProfiler* [35]. This method takes as input a list of DEGs and a list of all genes that are sufficiently expressed in the experiment (as determined by the filtering method of *edgeR* Table 1). The ORA method tests if a given gene set (a KEGG pathway or a GO term) is significantly over-represented in the input list of DEGs, relative to the background genes.

The second method, GSEA, was performed using the Python package *GSEApy* [36]. Similarly to ORA, this method also takes as input a list of all sufficiently expressed genes, albeit without an additional list of DEGs. Instead, the input genes are ranked by a metric chosen by the user, such as fold changes or p-values. The method then tests whether a given gene set is enriched at the extreme ends of the ranked gene list. Compared to ORA, this approach has two advantages. First, no arbitrary thresholds (for example, on FDR or log_2_ FC) are required to define any DEGs. Second, the chosen ranking metric incorporates additional, gene-level information into the analysis, assigning different weights to individual genes based on the strength of differential expression. By contrast, ORA relies on a dichotomization of the data into DEGs and non-DEGs, disregarding differences between individual genes in either of the thus-introduced groups.

For GSEA, we used two distinct ranking metrics. First, we ranked the genes by log_2_ FC, as this is a common approach and allows us to re-use the fold change values already computed by *edgeR*. (Results obtained using *DESeq2* fold change estimates are similar, as the fold change estimates between the tools differ little.) Second, we ranked the genes by the absolute value of the signal-to-noise ratio (|S2N|), as this metric was found to be among the best-performing metrics in a systematic comparison by Zyla et al. [28]. The absolute signal-to-noise ratio is defined as

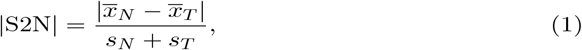

where the numerator quantifies the difference in mean read counts between normal and tumor tissue samples for a specific gene, and the denominator quantifies the sum of standard deviations for the same counts. Before the calculation, we normalized the counts using the *DESeq2* function *estimateSizeFactors* to account for differences in library sizes. Additionally, from the full gene set libraries, we dropped all gene sets with fewer than 15 or more than 500 genes, which is the default setting for *GSEApy* and close to the default setting for *clusterProfiler*. To make the results comparable, we took the intersection of gene sets tested by these two methods, leaving us with 191 KEGG terms and 1’390 GO terms as our final search space. To determine the significance of the gene sets, we again used a 5% threshold on the Benjamini-Hochberg adjusted p-values to control the FDR.

### 5.4 Experiment replicability

We proceed by introducing a fundamental performance metric used throughout the rest of this paper. The goal is to measure the level of mutual agreement between the results obtained from individual simulated experiments with the same cohort size and cancer type (Fig. 7). For this, we designate the set of analysis results as *S*_*i*_, where *S* can be a set of DEGs, a set of enriched GO terms, or a set of enriched KEGG pathways, and *i* ∈ {1, 2, …, 100} indexes the experiment. We then use the Jaccard index (intersection over union) to define the inter-experiment replicability of results from two simulated experiments,

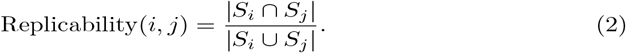

This is a value between zero (no overlap of results) and one (perfect agreement). If either *S* is empty, we define the replicability to be 0. We report the median experiment replicability over all 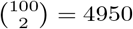 distinct pairs (*i, j*).

It should be noted that replicability is non-zero, on average, also when the analysis results (DEGs, enriched terms, or enriched pathways) are chosen at random (the null model). Therefore, as a benchmark, we introduce the expected replicability under the null model as

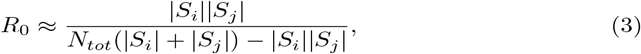

where *N*_*tot*_ is the total number of tested genes or terms/pathways. Rather than calculating this value for every pair (*i, j*), we calculated the median null replicability by setting |*S*_*i*_| = |*S*_*j*_| = median{ |*S*_*i*_| | *i* ∈ 1, 2, …, 100 }.

We continue with a derivation of Eq. (3). Given two random subsets *A* and *B* with fixed sizes 0 *<* |*A*|, |*B*| ≤ |*N* | drawn independently and uniformly without replacement from a parent set *N*, we want to know the expected value of the intersection of *A* and *B*. This problem is equivalent to determining the number of successes *k* one can expect in *n* draws without replacement from a set of size |*N* |, where *K* of its elements are considered successes and the remaining elements are considered failures. Thus reformulated, the problem defines the well-known hypergeometric distribution with mean *E*[*k*] = *nK/*|*N* |. Without loss of generality, we set |*A*| = *n* and |*B*| = *K* to find

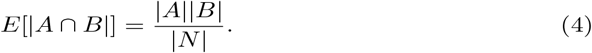

The expected value for the union of *A* and *B* is then simply

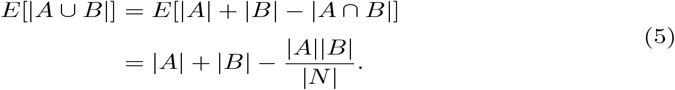

Finally, the expected value for the Jaccard index can be approximated by

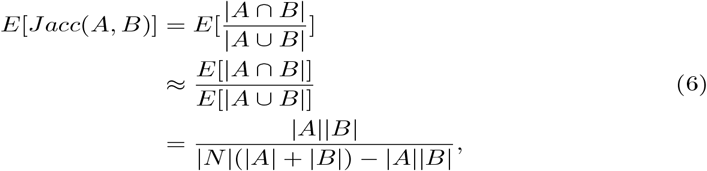

which is equivalent to our approximation of *R*_0_ in Eq. (3). In the second line we have used a first-order Taylor expansion around the expected values of the numerator and denominator. Simulations show that the error of this approximation rapidly becomes negligible with increasing set sizes. This further justifies the use of this approximation for our purposes, where sets of DEGs typically range from a few hundred to several thousands of genes.

### 5.5 Ground truth and binary classification metrics

An analysis pipeline for a given fold change thresholding method (zero, non-zero, post hoc) consists of using *edgeR* (QLF and LRT) or *DESeq2* (Wald) for differential expression analysis and fold change estimation, followed by ORA or GSEA for enrichment analysis. In addition to measuring experiment replicability, we defined pipeline-specific ground truths by running the respective pipeline on the full data set with all patients of the respective cancer type (Table 1). For DEGs, the final ground truth was obtained by taking the intersection of DEGs from all three statistical tests for differential expression analysis (QLF, LRT, Wald) for a given fold change threshold. These ground-truth DEGs were in turn used to define the ground truth of enriched gene sets for ORA. The ground truth genes for all seven cancer types with additional information are available in Section 5.9 (Availability of data and materials).

After having determined the ground truth for both DEGs and enriched gene sets, we calculated two classical binary classification metrics: precision and recall (also known as positive predictive value and sensitivity, respectively). In this way, we quantify how well the results from small cohort sizes approximate the results obtained from much larger cohort sizes. Additionally, we calculated the Matthew’s Correlation Coefficient (MCC) [37], which is a balanced performance metric that incorporates all four categories of the confusion matrix and ranges from −1 (worst) to +1 (best). The used metrics are defined as

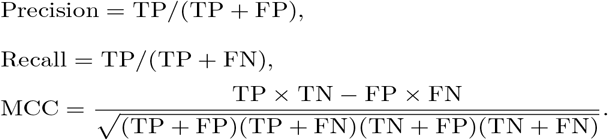

Similarly to replicability, expected values for these metrics can be computed under the null model of random gene selection. Given a set of genes *N*, a non-empty subset of ground truth genes *G* ⊂ *N*, and a uniformly drawn non-empty subset of significant genes *S* ⊂ *N*, it is straightforward to calculate expected values for the binary classification metrics (precision, recall, MCC). Let *s* ∈ *S*, then

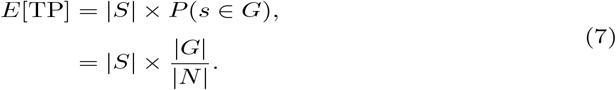

The expected values for FP, TN, FN can be obtained directly from *E*[TP] and |*G*|, |*S*|, |*N* |. The metrics are then readily obtained as

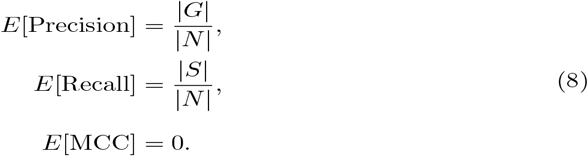

### 5.6 Outlier removal

In addition to testing different analysis pipelines, we were interested in testing whether outlier removal can improve the robustness of the analysis results. To this end, we tested two different methods to detect outlier patients: a recently introduced robust principal component analysis (rPCA) method [22] and a custom algorithm based on jackknife resampling.

The rPCA method uses robust statistics to obtain principal components that are not substantially influenced by outliers. Additionally, this method allows for the detection of outlier samples based on objective criteria, rather than subjectively, through the visual inspection of the classical PCA plot. For our study, we used the *PcaHubert* method from the *rrcov* R package with *k* = 2 principal components to flag outlier patients in a given cohort. The choice of *k* = 2 is a typical value and allows visual validation of the results [22].

Additionally, we implemented a custom algorithm to detect outlier patients based on jackknife resampling. We designate the method *ReBoost*, as it is designed to maximize (boost) the recall of DEGs at a fixed significance level. It works as follows: Given a cohort with *N* patients, we first determine the initial number of DEGs using the *edgeR* QLF method. Next, we determine the number of DEGs in each subcohort of size *N* − 1 obtained by omitting one patient. A patient is marked as an outlier if the number of DEGs increases upon removal of the patient relative to the initial number of DEGs. After removing all marked outlier patients, the procedure is repeated iteratively until no more patients can be removed.

## Supplementary information

### Declarations

#### 5.7 Ethics approval and consent to participate

Not applicable.

#### 5.8 Consent for publication

Not applicable.

#### 5.9 Availability of data and materials

The datasets supporting the conclusions of this article are available in the GDC repository, [https://portal.gdc.cancer.gov/]. A persistent Git repository with Python scripts and notebooks for downloading the raw data and performing the analysis can be found on Zenodo [https://doi.org/10.5281/zenodo.8333519]. The repository also includes processed (aggregated) datasets which we used to generate the figures, as well as lists of ground truth DEGs for each cancer type.

#### 5.11 Authors’ contributions

PD wrote the code, contributed to the design of the study, and analyzed the results. MM obtained funding, planned, and supervised the study. Both authors wrote, read, and approved the manuscript.

## 5.12 Acknowledgements

Calculations were performed on UBELIX (http://www.id.unibe.ch/hpc), the HPC cluster at the University of Bern.

